# Mesenchymal Stromal Cells Donate Mitochondria to Articular Chondrocytes Exposed to Mitochondrial, Environmental, and Mechanical Stress

**DOI:** 10.1101/2022.05.12.491696

**Authors:** Megan J. Fahey, Maureen P. Bennett, Matthew Thomas, Irene Vivancos-Koopman, Lindsay Browning, Lawrence J. Bonassar, Michelle L. Delco

## Abstract

Avascular soft tissues of the skeletal system, including articular cartilage, have limited healing capacity, in part due to their low metabolic activity. No drugs are available that can prevent or slow the development of osteoarthritis (OA) after joint injury. Therefore, mesenchymal stromal cell (MSC)-based regenerative therapies are increasingly common in the treatment of OA, but questions regarding their clinical efficacy and mechanisms of action remain unanswered. Our group recently reported that mitochondrial dysfunction is one of the earliest responses of cartilage to injury, resulting in chondrocyte death, extracellular matrix degeneration, and ultimately OA. MSCs have been found to rescue injured cells and improve healing by donating healthy mitochondria in highly metabolic tissues, but mitochondrial transfer has not been investigated in cartilage. Here, we demonstrate that MSCs transfer mitochondria to stressed chondrocytes in cell culture and in injured cartilage tissue. Conditions known induce chondrocyte mitochondrial dysfunction, including stimulation with rotenone/antimycin and hyperoxia, increased transfer. Stressed chondrocytes increased expression of genes related to inflammation and senescence, further supporting the link between mitochondrial dysfunction and transfer. MSC-chondrocyte mitochondrial transfer was blocked by non-specific and specific (connexin-43) gap-junction inhibition. When MSCs were exposed to mechanically injured cartilage they localized to areas of matrix damage and extended cellular processes deep into microcracks, delivering mitochondria to chondrocytes. This work provides insights into the chemical, environmental, and mechanical conditions that can elicit MSC-chondrocyte mitochondrial transfer in vitro and in situ, and our findings suggest a new potential role for MSC-based therapeutics after cartilage injury.

**Significance Statement:** Recent evidence suggests that although articular cartilage is avascular and relatively metabolically quiescent, acute injury induces chondrocyte mitochondrial dysfunction, driving cartilage degradation and OA. We present the first evidence that MSCs donate mitochondria to articular chondrocytes undergoing mitochondrial dysfunction in vitro and in situ. These findings support a new role for MSCs in the context of cartilage injury and OA, and intercellular mitochondrial transfer may represent a new biological approach to augment mitochondrial capacity in injured chondrocytes. This work establishes multiple experimental models to study MSC mitochondrial donation for the treatment of OA and related degenerative diseases of avascular orthopedic tissues.

## Introduction

ATP generated by mitochondrial (MT) oxidative phosphorylation is critical for cell survival and repair, and MT are the main source of reactive oxygen species in most tissues. Therefore, it is unsurprising that MT dysfunction is a fundamental pathology underlying many degenerative diseases of highly metabolic tissues, including chronic obstructive pulmonary disease (1), degenerative retinopathy (2), ischemic cardiomyopathy (3), and Parkinson’s (4).

Amongst diverse tissue types, there is a spectrum of reliance on MT metabolism; cardiomyocytes heavily utilize MT oxidative phosphorylation to sustain high energy demand, while chondrocytes, the sole cell type in articular cartilage, are highly glycolytic, producing only 10-25% of their ATP by MT respiration (5, 6). Further, articular cartilage is avascular, and chondrocytes are adapted to extreme environmental conditions, including low nutrient availability and oxygen concentrations of ∼5% (7). Cartilage contains at least an order of magnitude fewer cells per tissue volume and only ∼5% of the MT volume per cell (5) compared to liver. Taken together, this evidence would seem to minimize the importance of MT in cartilage health and disease, and may explain why the role of MT dysfunction in the initiation and pathogenesis of osteoarthritis (OA) has not been well studied (8). Recent work by our group and others revealed that MT dysfunction is one of the earliest responses of cartilage to injury, resulting in cell death, degeneration of the extracellular matrix, and ultimately post-traumatic OA (9, 10). We also found that early pharmacological intervention using the MT protective peptide SS-31 after cartilage injury reduced chondrocyte death and prevented cartilage degeneration (11–14). While these studies provide rationale for targeting MT function to prevent OA and similar degenerative diseases of avascular tissues, strategies for restoring MT function to improve tissue healing have not been developed.

Mesenchymal stromal cells (MSCs) are being widely investigated as regenerative therapies, and increasing evidence suggests MSCs can relieve symptoms of OA including pain and joint dysfunction, as well as preserve cartilage (15, 16). While the mechanisms governing the beneficial effects of implanted cells remain unclear, contemporary evidence indicates that MSCs act mainly by modulating the joint environment via their secreted products, which inhibit catabolic signaling pathways and promote anabolic responses (17).

Intriguingly, recent evidence suggests an alternate paradigm; MSCs can rescue injured cells undergoing MT dysfunction by donating functional MT. This intercellular MT transfer restores bioenergetics and preserves viability of recipient cells under metabolic stress (18) In vitro, MT transfer was evident within 12 hours of co-culture between MSCs and vascular endothelial cells subjected to ischemia-reperfusion injury (19). In an in vivo model of acute lung injury from endotoxin installation into the airways of mice, MT transfer from MSCs to alveolar epithelial cells increased ATP concentration (20). These and other studies of MT transfer have involved oxidative cell types, such as cardiomyocytes (3) and neurons (21). While chondrocytes are not considered oxidative, the role MT dysfunction in OA supports investigating MT transfer in cartilage. Wang et al. (2020) reported evidence of MT transfer to chondrocytes in cell culture using fluorescent imaging and that MT transfer is associated with improvement of MT function(22); however, MT transfer was not confirmed or quantified, and the mechanisms of transfer have not been investigated. Although the specific mechanisms of intercellular MT transfer are still largely unknown, several processes have been implicated in MT transfer between MSCs and dissimilar cell types, including tunneling nanotubule formation, gap junction signaling, extracellular vesicle communication, and cell-cell fusion events (23–25). Importantly, the in vivo model of acute lung injury found that MSCs transferred MT by integrating with the surrounding tissue in a connexin 43 (Cx43)-dependent manner (20). Cx43 is the protein monomer of gap junctions that connect adjacent cells and is known to be expressed by chondrocytes (26). Notably, chondrocyte Cx43 expression is increased in OA (27) and when stressed with IL-1ß (28).

Current literature suggests that implanted MSCs can be recruited to damaged tissues by cells undergoing metabolic stress and establish gap junctions to donate MT. However, this has not been investigated in cartilage. Therefore, our goal was to develop model systems to study MT transfer from MSCs to chondrocytes, study factors that increase MT transfer, and begin to identify the mechanisms involved. We hypothesized that MT transfer from MSCs to chondrocytes would occur in vitro and in situ, factors that inhibit chondrocyte MT function would increase MT transfer from MSCs, and that inhibiting gap junctions, specifically Cx43, would decrease transfer. Identifying mechanisms that mediate MT transfer in cartilage is the first step toward developing approaches to exploit this biological phenomenon to improve healing in avascular orthopedic tissues.

## Results

### MSCs transfer MT to chondrocytes in vitro

To determine if MT transfer occurs between MSCs and chondrocytes, a primary cell co-culture model was used; equine articular chondrocytes and MSCs were harvested, cultured, and stained with Calcein AM or MitoTracker Deep Red and Hoescht, respectively. Chondrocytes were stressed with MT-specific inhibitors rotenone and antimycin (Rot/A) then co-cultured with MSCs for up to 8 hours at physiologic oxygen concentration for chondrocytes (5% O_2_; Fig 1A). Flow cytometry was used to quantify MT transfer events; separately cultured chondrocytes and MSCs were used to set up quadrant gates that identified chondrocytes by green fluorescence and MSCs by red fluorescence (Fig 1B; top plot). This gating strategy was used on co-cultured cells, which allowed us to quantify the upper right-hand quadrant of cells positive for both stains (red^+^/green^+^ cells), identifying what we considered transfer events (Fig 1B; bottom plot). The percentage of red^+^/green^+^ cells increased with duration of co-culture up to 8 hours (Fig 1C). Confocal microscopy confirmed that stimulated chondrocytes gained red MT fluorescence, and MSCs gained green fluorescence, indicative of intercellular MT transfer. Several modes of transfer were visualized, as previously described for other cell types (18) including apparent microvesicle transfer (Fig 1Di), with and without evidence of cell-cell contact (Fig 1Dii), tunneling nanotubule-mediated filopodial transfer (Fig 1Diii), and cell fusion events (Fig 1Div,v). There are limitations associated with live cell staining techniques including MT toxicity from stains, short-term fluorescence, and rapid photobleaching. We also observed the indirect staining of equine chondrocytes with Hoescht from co-culture with the Hoescht-stained MSCs (Fig 1D). Given these limitations, we further investigated intercellular MT transfer using transgenic mouse strains expressing endogenous fluorescent proteins.

**Figure 1.**
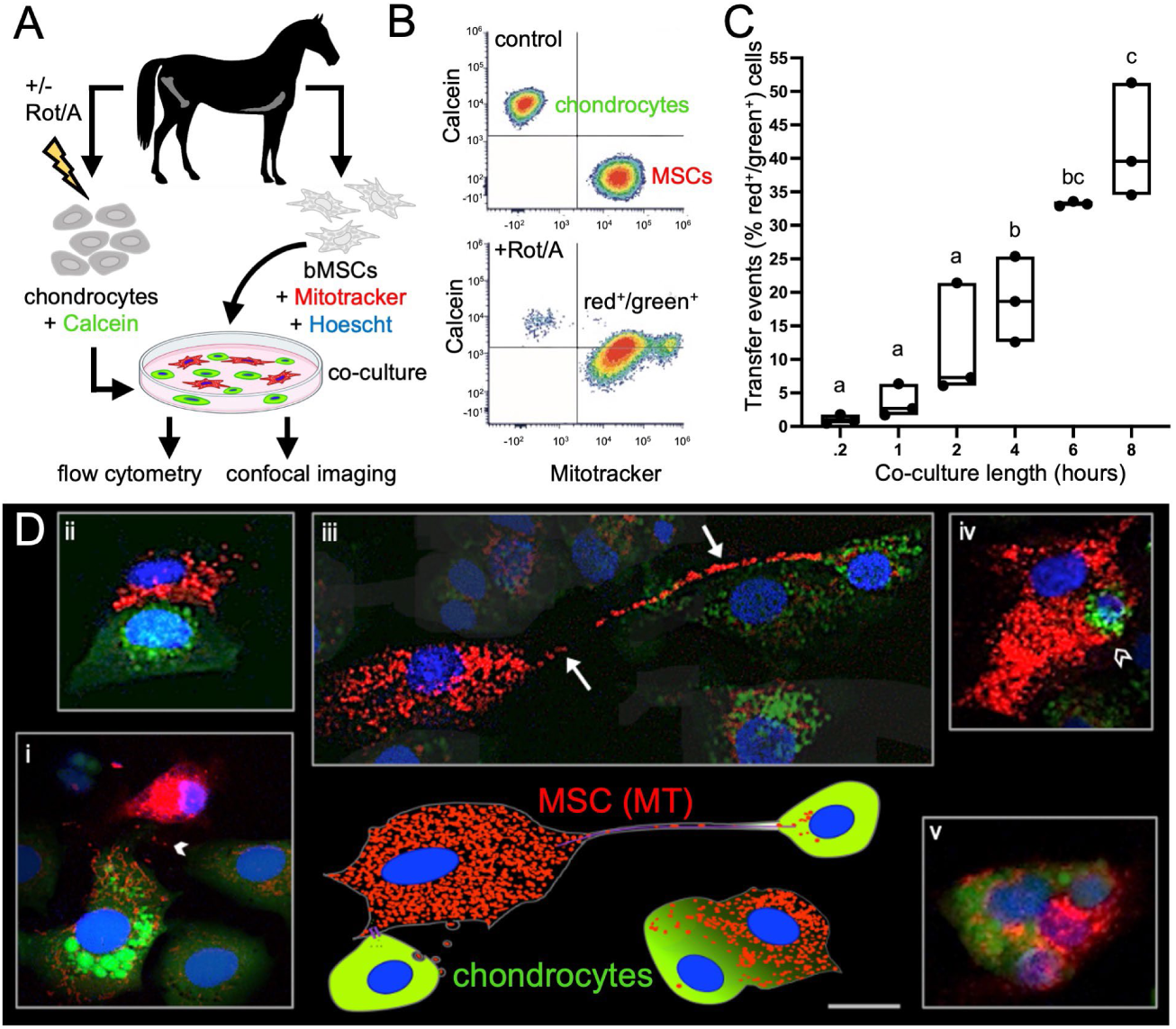
Equine mesenchymal stem cells (MSCs) transfer mitochondria (MT) to chondrocytes over time and via several modes. A) Experimental design B) Representative flow cytometry plots. Top plot shows results of separately cultured equine chondrocytes stained with Calcein AM (4mM) and MSCs stained with MitoTracker Deep Red (200nM) used to set quadrant gates; bottom plot shows red^+^/green^+^ chondrocytes (upper right-hand quadrant) after rotenone and antimycin (Rot/A; 0.5 μM/0.5 μM). stimulation and co-culture with MSCs. (C) MT transfer (% red^+^/green^+^ cells) increases with length of co-culture, n = 3. (D) Schematic (bottom center) depicting observed modes of MT transfer from MSCs (red MT, blue nuclei) to chondrocytes (green cytoplasm, blue nuclei) alongside corresponding confocal microscopy images (i-v), taken up to 8 hours after initiation of co-culture, stained as above with the addition of Hoechst (5 mg/L, blue) to MSCs before co-culture; representative images of (i) likely gap junction-mediated microvesicle (arrowhead) transfer, (ii) close cellular contact without apparent transfer demonstrating the broad range of observed cell-cell interactions, (iii) possible tunneling nanotubule-mediated filopodial (arrows) transfer, (iv) apparent phagocytosis-like event (open arrowhead), (v) cell fusion event. Note that in (i) and (iii), contrast in surrounding cells was decreased 10-20% to highlight the mentioned interaction; Bar ≅ 10µm.

Articular chondrocytes were harvested and cultured from UBC mCherry mice, with ubiquitous red fluorescent cytoplasmic protein expression, and MSCs were isolated and expanded from PhAM mitoDendra2 mice, which express a MT-targeted green fluorescent protein. Co-culture of these cell types allowed flow cytometry and confocal imaging to confirm intercellular MT transfer (Fig 2A). Longitudinal visualization of cellular interactions over 9.5 hours of co-culture revealed MSCs shedding green MT (Fig 2Ei; white arrowheads) and localization of green MT in red chondrocytes at 7.5 (Fig 2Eii) and 9.5 hours (Fig 2Eiii) of co-culture.

**Figure 2.**
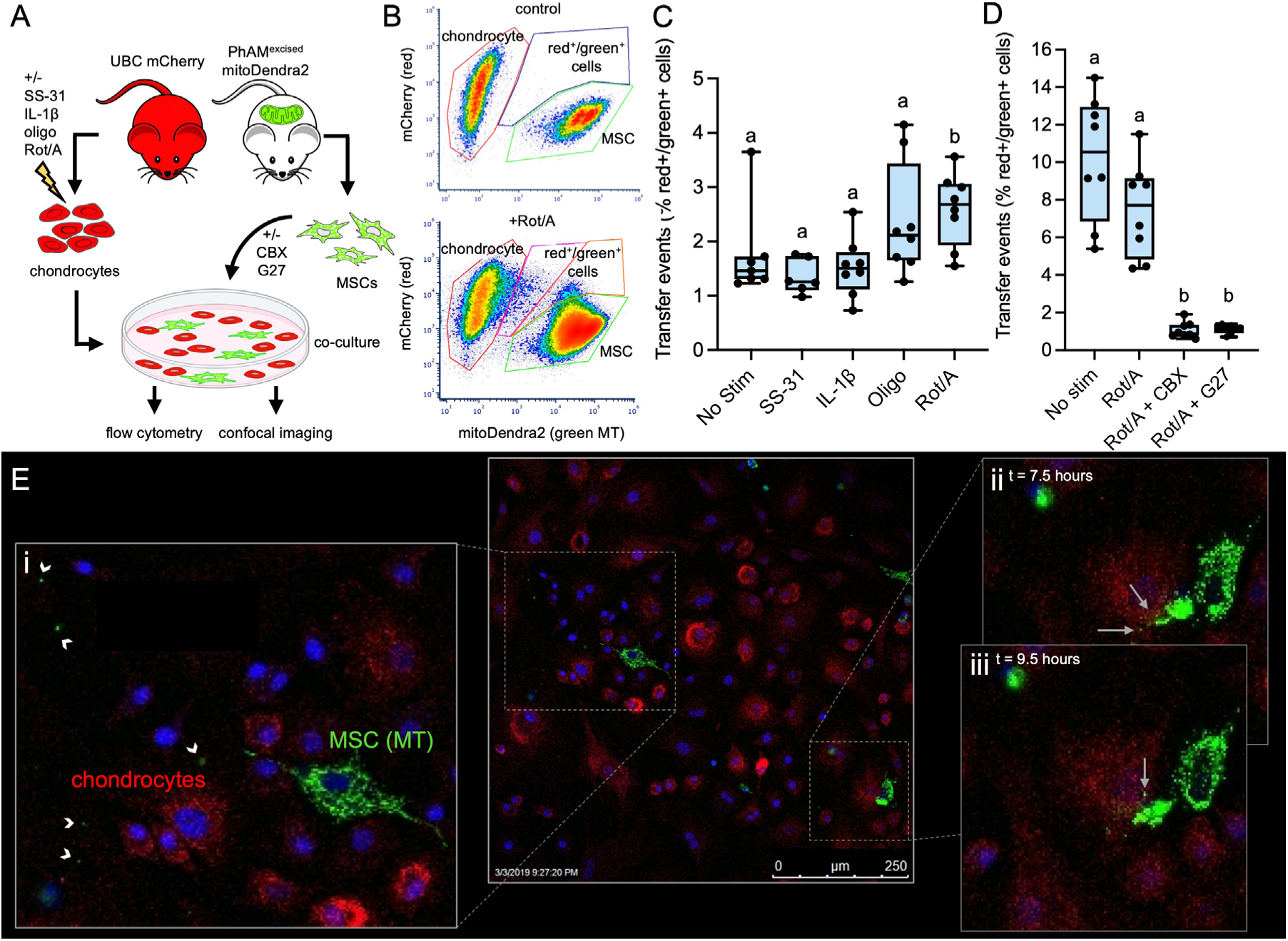
Murine mesenchymal stem cells (MSCs) donate mitochondria (MT) to chondrocytes and transfer is increased by MT-specific stimulants and decreased with gap-junction inhibitors. (A) Experimental design (B) Representative flow cytometry plots. Top plot shows results of separately cultured mitoDendra2 MSCs and mCherry chondrocytes used to set gates; bottom plot shows chondrocytes co-cultured with MSCs after stimulation with rotenone and antimycin (Rot/A; 0.5 μM/0.5 μM). (C) MT transfer events (% red^+^/green^+^ cells) from MSCs to chondrocytes after 12-hour co-culture is significantly higher when chondrocytes are stimulated with the MT-specific stressor Rot/A under physiologic culture conditions (5% O_2_, 0.45 g/L glucose), n = 4. (D) Both carboxonolone disodium (CBX; 100 μM) and gap 27 (G27; 100 μM) significantly inhibited MT transfer events. MSCs were treated with these gap junction inhibitors during 24-hour co-culture, n = 4 (E) Live confocal microscopy of MT transfer. MSCs were stained with Hoechst (5 mg/L) for visualization of nuclei. (i) Apparent MSC microvesicles containing MT (white arrowheads); image acquired 4.5 hours after initiation of co-culture, (ii) same position 3 hours later, MSC MT transferred to chondrocyte (white arrows) (iii) same position 5 hours later. Co-culture was conducted in growth medium on Leica SP5 inverted confocal microscope outfitted with temperature and humidity-controlled chamber, using a 40x objective lens. Scale bar = 250 μm.

### Inhibition of MT respiration affects MT transfer events

We investigated if MT dysfunction in chondrocytes is associated with increased intercellular MT transfer in our transgenic murine model. Chondrocytes were cultured under physiologic oxygen and glucose concentrations (5% O_2_ and 0.45 g/L glucose), stimulated with either a general inflammatory stimulus (IL-1ß) or MT-specific inhibitors (Rot/A or oligomycin (Oligo)) prior to co-culture with MSCs, and evaluated using flow cytometry. Because 2-dimensional culture alone is a known stressor in chondrocytes, as an additional negative control chondrocytes were treated with SS-31, a MT-protective peptide that improves ATP turnover and prevents proton leak in chondrocytes (11). Gates for red^+^ and green^+^ cells were created based on non-co-cultured controls (Fig 2B; top plot). These gates were used to quantify red^+^/green^+^ cells after treatments and co-culture with MSCs (Fig 2B; bottom plot). We found that inducing MT dysfunction by inhibiting the electron transport chain with Rot/A increased the percentage of red^+^/green^+^ cells compared to the other stimulants and non-stimulated control (Fig 2C). The ratio of chondrocytes to MSC in this experiment was 10:1. When the ratio was altered to 2:1, Rot/A did not affect transfer events but the percentage of red^+^/green^+^ cells was higher in both non-stimulated and Rot/A stimulated chondrocytes (Fig 2D). Pre-treatment with SS-31 limited the rate of MT transfer to <2% but this was not significantly different from the non-stimulated control group (Fig 2C).

### Gap junction inhibition prevents MSC-chondrocyte MT transfer

Gap junctions allow physical interaction and direct signaling between dissimilar cell types, and have been implicated in MSC MT donation (20, 29). We investigated this mechanism in our co-culture system by treating murine chondrocytes and MSCs with gap junction inhibitors and assessing their effect on the percentage of red^+^/green^+^ cells. Carbenoxolone disodium (CBX) is a derivative of 18-alpha-glycyrrhetinic acid and a non-specific inhibitor of gap junction electrical coupling (30). Treatment with CBX reduced the percentage of red^+^/green^+^ cells by 7-fold (Fig 2D). A more specific inhibitor, Gap 27 (G27), was used to reversibly inhibit connexin 43 (Cx43)-mediated cell-cell communication (31), which prevented MT transfer to a similar extent as CBX (Fig 2D).

### Culture conditions affect MSC-chondrocyte MT transfer

Because homeostatic chondrocytes are adapted to low oxygen and nutrients conditions in vivo, we evaluated the effects of environmental culture conditions on MT transfer. Murine chondrocytes were isolated and cultured at physioxia (5% O_2_) or hyperoxia (21% O_2_), and euglycemia (0.45 g/L) or hyperglycemia (1 g/L), then stressed with Rot/A or IL-1ß. The percentage of red^+^/green^+^ cells was determined by flow cytometry as previously described. Non-physiologic culture conditions alone, and their interaction with stimulation, affected MT transfer events (Figure 3; p<0.0001).

**Figure 3.**
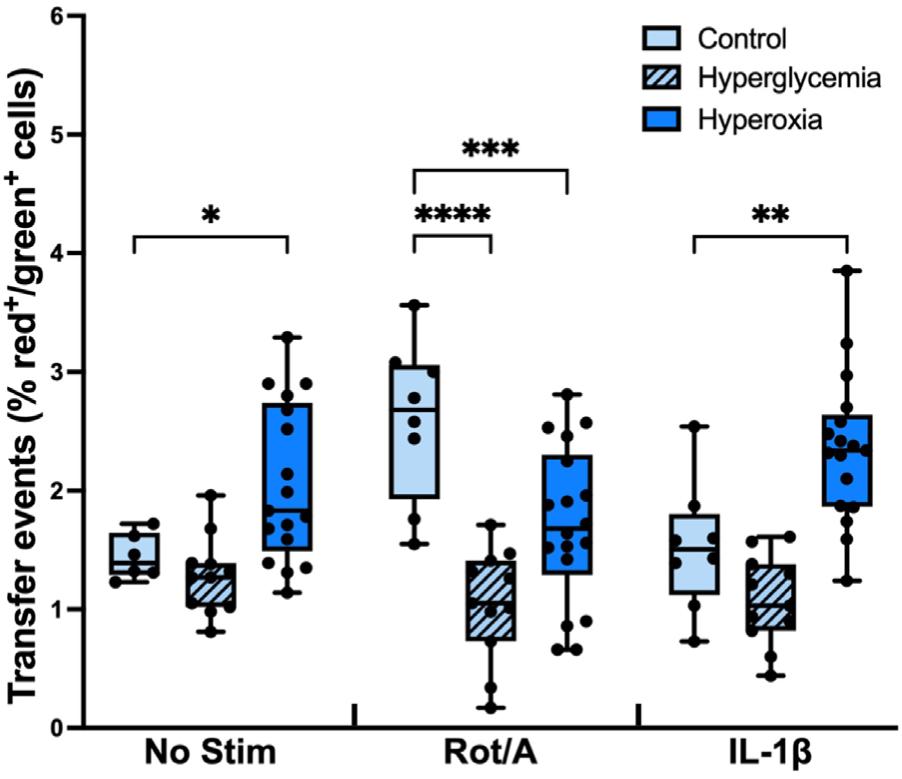
MSC-chondrocyte MT transfer is affected by non-physiologic culture conditions. Murine mCherry chondrocytes (red) and PhAM mitoDendra2 MSCs (green MT) were co-cultured in hyperoxia or hyperglycemia for 12 hours after stimulation with or without Rot/A or IL-1β. MT transfer events (% red^+^/green^+^ cells) were quantified by flow cytometry. Hyperoxia increases transfer events in non-stimulated and IL-1β-stimulated chondrocytes (P < 0.05). Hyperoxia and hyperglycemia decreased transfer events when chondrocytes were stimulated with Rot/A (P < 0.0007). n = 4, N = 107.

Hyperoxia increased MT transfer in both non-stimulated and IL-1ß-stimulated chondrocytes (p<0.0017). Hyperglycemia caused MT transfer to decrease in Rot/A chondrocytes relative to Rot/A chondrocytes in physiologic conditions. When chondrocytes were non-stimulated or stimulated with IL-1ß, hyperglycemia did not affect transfer.

### Hyperoxia increases chondrocyte expression of gap junction, inflammation and senescence associated genes

To begin to understand the link between increased MT transfer and MT dysfunction in chondrocytes, we developed a custom quantitative PCR panel of relevant genes involved in chondrocyte metabolism and OA. We found that expression of gap junction alpha 1 (GJA1), the gene that encodes the Cx43 protein, was increased in hyperoxia (Fig 4A). The senescence associated markers, cyclin-dependent kinase inhibitor 2A (CDKN2A) and tissue inhibitor of metalloproteinases 1 (TIMP1), were also increased in hyperoxia (Fig 4B,C). In addition, hyperoxia caused chondrocytes to increase the expression of the antioxidant enzyme, SOD2 (Fig 4D).

**Figure 4.**
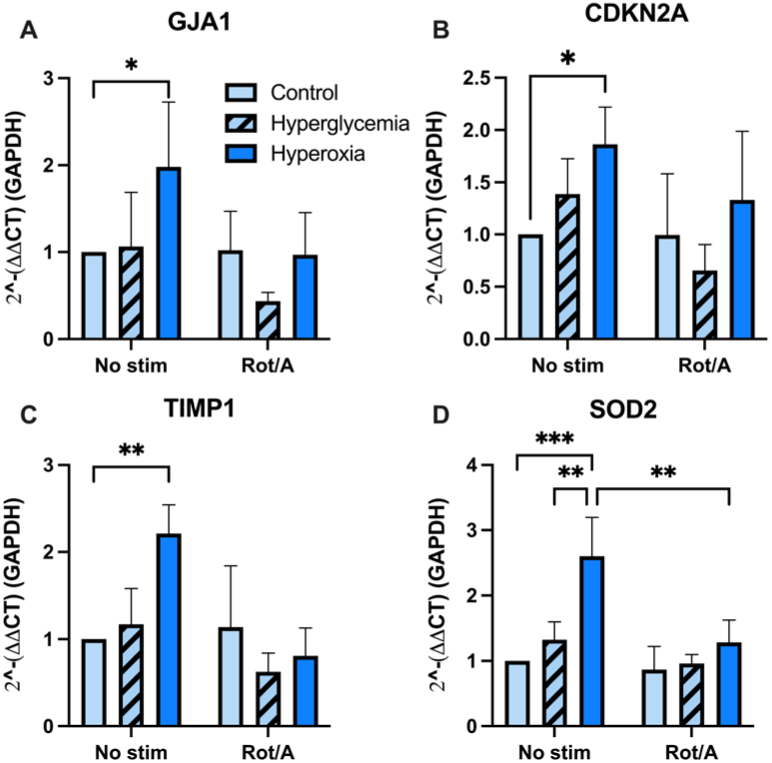
Chondrocyte gene expression changes in hyperoxia and with rotenone/antimycin inhibition. A) Gap junction protein alpha 1 (GJA1) expression increased in non-stimulated chondrocytes cultured in hyperoxia, n=4. B) Cycling dependent kinase inhibitor 2A (CDKN2A) expression increased in hyperoxia, n=4. C) Tissue metallopeptidase inhibitor 1 (TIMP1) expression increased in non-stimulated chondrocytes cultured in hyperoxia, n=4. D) Superoxide dismutase 2 (SOD2) expression increased in chondrocytes cultured in hyperoxia, n=4.

### MSCs transfer MT to chondrocytes in injured cartilage explants

To assess whether MSCs can deliver MT to chondrocytes embedded in cartilage tissue *in situ*, we utilized an established model of cartilage injury, in which cartilage explants are impacted at loading rates capable of inducing extracellular matrix damage and MT dysfunction (9, 32, 33). Bovine cartilage explants were injured and stained with CFDA, a green cellular dye, then incubated with bovine MSCs transduced with mCherry-mito, a MT-targeted red fluorescent protein, for 4 days (Fig 5A). Confocal imaging revealed that MSCs adhered to the articular surface and localized to areas of matrix damage in injured explants (Fig 5C). In contrast, MSCs were rarely identified on the surface of uninjured cartilage (Fig 5B). Cross-sectional confocal imaging of injured explants (Fig 5D) revealed that MSCs extended into cracks in the cartilage matrix to depths of more than 100 µm from the articular surface. MSC MT co-localized with chondrocytes near microcracks and in direct contact with MSCs. Deeper in the tissue, MSC MT were identified within chondrocytes at >50um from cracks, with no evidence of physical interaction between chondrocytes and MSCs.

**Figure 5.**
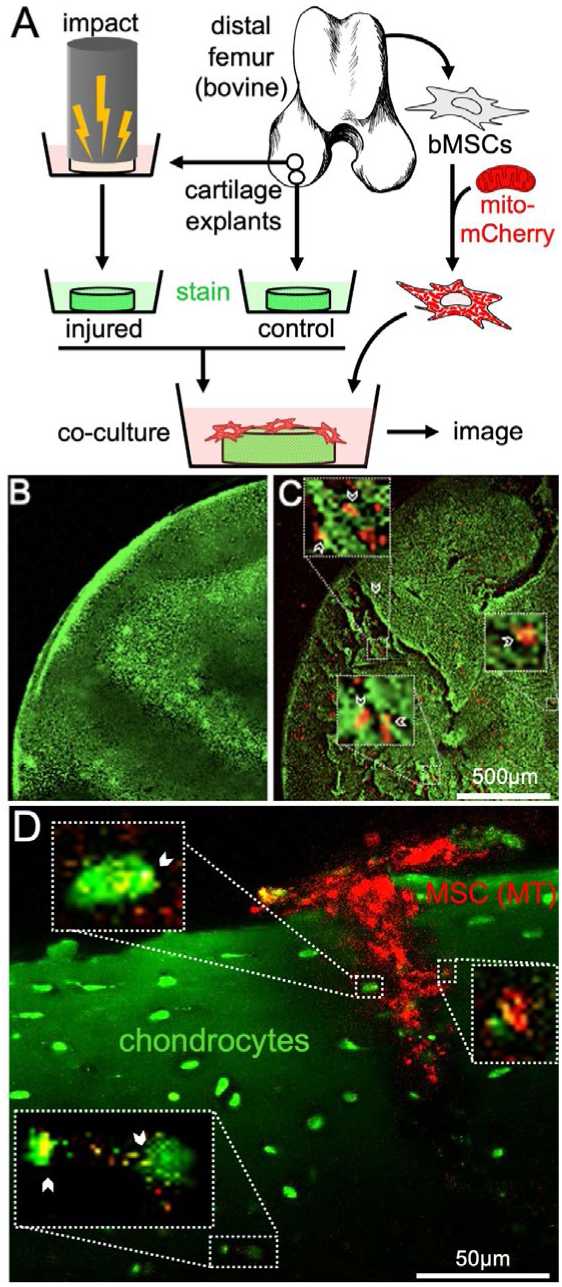
Bovine mesenchymal stem cells (MSCs) home to damaged cartilage and transfer mitochondria (MT) to chondrocytes in situ. A) Schematic depicting experimental design for cartilage explant injury. B) Representative image of unimpacted cartilage explant stained with CFDA (green) and seeded with lentiviral transduced mCherry-mito MSCs (red MT) compared to C) injured cartilage seeded with MSCs and cultured for 4 days. MSCs localized to damaged areas on injured cartilage, while MSCs were not present on the surface of uninjured cartilage. (D) High magnification images of MSC (red MT) localized deep within a crack in injured cartilage (articular surface is toward the top). MSC interactions with chondrocytes (green) are evident in several locations (white arrowheads) and MT transfer is evident in areas of red^+^/green^+^ colocalization (yellow) close to the crack (top insets) as well as remote and deep to the crack (bottom inset).

## Discussion

This study provides evidence of intercellular MT transfer from bone marrow derived MSCs to articular chondrocytes in cell culture and in injured cartilage tissue. The phenomenon of MT donation by MSCs has primarily been described in energy-expensive tissues, such as heart and brain, which heavily utilize MT oxidative phosphorylation for ATP production (3, 21). Although healthy cartilage derives the majority of its energy from glycolysis, MT function is critical to chondrocyte homeostasis and extracellular matrix (ECM) synthesis. The highly ordered structure of the ECM endows articular cartilage unique functional properties of cushioning and lubricating joints. Notably, mice with impaired MT electron transport chain proteins had altered ECM composition, resulting in increased cartilage stiffness (34). MT are highly plastic organelles that undergo morphological and functional changes in response to synthetic and bioenergetic demands (35). However, recent work revealed that in chondrocytes, MT polarity and oxygen consumption rate per viable cell decreases in the hours following mechanical injury (9, 36). This suggests that with few MT, the ability of chondrocytes to upregulate ATP production and biosynthetic capacity in the critical period following injury is limited. While chondrocytes can increase MT content via biogenesis, this requires time (hours-days) and an initial energy investment. Taken together, this evidence suggests that while MT diseases and MT-targeted therapies are generally associated with metabolically active, highly oxidative tissues, cells with minimal spare respiratory capacity such as chondrocytes may recruit MT donation as a rescue mechanism during times acute of stress. Further studies are necessary to investigate the functional impact of MT transfer on cartilage.

MT dysfunction has recently been identified as a key subcellular event in the initiation and early pathogenesis of OA (37, 38). Although the mechanisms are unclear, MT dysfunction in recipient cells appears to be a prerequisite for the initiation of MT donation by MSCs in other cell types (18), and our findings support this in chondrocytes. We investigated various stimuli known to induce MT dysfunction. IL-1ß is a well-established in vitro model of OA, and this ‘master’ pro-inflammatory cytokine has been found to play a central role in disease pathogenesis (39). Further, inflammatory cytokines including IL-1ß and TNF-alpha induce chondrocyte MT dysfunction by indirectly inhibiting the activity of MT Complex I (40). While IL-1ß did not affect MT transfer in our murine co-culture system, a more specific MT inhibitor, Rot/A increased transfer events. Rot/A inhibits MT complexes I and III, respectively. By inhibiting the flow of electrons through the electron transport chain, Rot/A prevents ATP production while also increasing reactive oxygen species (ROS) due to incomplete transfer of electrons to oxygen in the final step of oxidative phosphorylation (41). Our findings suggest that bioenergetic and/or oxidative stress in chondrocytes may induce MT transfer. Conversely, we found that Oligo, which blocks ATP synthase, did not increase transfer events. Since complexes I and III are the major sites of MT ROS production (5), these findings suggest oxidative stress may be more important than bioenergetic stress in eliciting intracellular transfer in this model.

Non-physiologic oxygen and nutrient conditions are known to affect chondrocyte metabolism. Of note, relative hyperoxia (21% O_2_) and hyperglycemia (1 g/L glucose) are considered standard conditions for culturing articular chondrocytes, but are not representative of the low nutrient availability and avascular nature of cartilage (5). Therefore, we compared transfer events to chondrocytes at physioxia (5% O_2_) versus relative hyperoxia (21% O_2_) and euglycemia (0.45 g/L glucose) versus relative hyperglycemia (1.0 g/L glucose). As expected, we found that hyperoxia caused an increase in transfer events in both non-stimulated and IL-1ß-stimulated chondrocytes, supporting our hypothesis that hyperoxia-induced MT dysfunction triggers MT transfer. Under these same conditions, the expression of the MT-specific antioxidant SOD2, was increased. This is consistent with literature demonstrating that chondrocytes cultured in hyperoxia experience oxidative stress due to increased superoxide production (5). Taken together, these data suggest oxidative stress caused by hyperoxia can induce MT transfer. Interestingly, chronic MT dysfunction in end-stage OA has been associated with decreased SOD2; OA chondrocytes from patients undergoing hip replacement surgery demonstrate SOD2 deficiency, which has been causally linked to oxidative damage mediated by MT-derived ROS (37, 38).

In further support of the hypothesis that MT-derived ROS triggers MT transfer from MSCs, chondrocytes increased expression of inflammation- and senescence-associated genes in hyperoxic culture; Increased TIMP1 expression, a regulator of metalloproteinases and ECM turnover, is indicative of an inflammatory phenotype in chondrocytes (42). We also found that CDKn2a, a marker of chondrocyte senescence (43) was increased in hyperoxia. These findings are in agreement with studies that reveal excess MT-derived ROS upregulate release of inflammatory cytokines including IL-1ß, cartilage ECM-degrading enzymes matrix metalloproteinase 1 and 3, and senescence associated β-galactosidase in part through activation of the NF-kB signaling pathway in chondrocytes (44–47).

In other tissue types, Cx43 and other gap junction proteins have been found to play a critical role in mediating intercellular MT transfer (18, 20). In an endotoxin model of acute lung injury, Cx43 was required for MSCs to attach to alveoli and generate Cx43-positive nanotubules and microvesicles, which resulted in MT transfer and increased intracellular ATP in recipient cells (20). We found that co-cultures treated with the Cx43-specifc inhibitor Gap27 had a 7-fold decrease in transfer events. Gap27 is a small, synthetic mimetic peptide that reversibly inhibits the channel function of Cx43 (31). This suggests that Cx43-based MSC-chondrocyte signaling is required for MT transfer in our model. Interestingly, Mayan et. al found that chondrocyte GJA1 (the gene encoding Cx43) expression and protein content was increased in OA cartilage (27). Here, we report that hyperoxic stress increases both MT transfer and GJA1 expression in chondrocytes. On confocal imaging of co-cultures, we observed several modes of transfer associated with connexin signaling in other tissues, including apparent filopodial and microvesicle-associated transfer. Further studies are necessary to elucidate specific mechanisms of MSC-chondrocyte MT transfer, and determine which processes predominate under specific conditions, including naturally occurring OA.

While MT transfer has been documented in several highly cellular tissues, a major potential barrier to MT transfer in articular cartilage is the low cellularity and high density of the extracellular matrix (ECM); The small pore size and highly charged nature of the cartilage ECM has been shown to inhibit transport of large particles such as antibodies and other investigational biologic therapies (48, 49). Therefore, in order to investigate the relevance of MSC-chondrocyte MT transfer in situ, we utilized a well-validated model of cartilage impact injury (9). Following injury, MSCs localized to areas of cartilage matrix damage, and donated MT to adjacent chondrocytes, confirmed by 3 dimensional colocalization of MSC MT fluorescent signal within chondrocytes. The striking frequency of direct MSC-chondrocyte interactions at sites of matrix damage suggests a signaling pathway, whereby MSCs home to and/or chondrocytes recruit MSCs to areas of tissue damage to initiate transfer. This is supported by literature that implicates ROS, TNFα, and NFκB signaling in triggering MT transfer from MSCs to other cell types (18). Further studies are necessary to investigate such mechanisms in cartilage.

Cross-sectional imaging of injured explants revealed evidence of close cellular interactions and MT transfer from MSCs to chondrocytes located immediately adjacent to microcracks in the cartilage ECM. We also identified recipient chondrocytes greater than 50 µm from crack interfaces. Previous studies have documented MT transfer via extracellular vesicles (18). As such, the finding of apparent non-contact MT transfer in cartilage may be a result of MSCs exporting MT in extracellular vesicles which are taken up by chondrocytes embedded within distant lacunae. This is supported by our in vitro longitudinal imaging studies, where MSCs appeared to shed MT into the extracellular environment. However, the highly charged ECM of cartilage may present a barrier to passive diffusion of these large, membrane-bound particles. Alternatively, it is possible that our in situ imaging studies failed to detect limited sites of MSC-chondrocyte contact. Recent evidence suggests that, contrary to conventional wisdom, chondrocytes are not isolated within lacunae but rather connected to one another by long, thin filapodial arms that extend through the ECM (50). In that study, Cx43-based gap junctions were found to be enriched at sites of filopodial contact. Given our findings implicating Cx43 in MSC-chondrocyte MT transfer, further investigation into gap junction signaling between MSCs and in situ chondrocytes is warranted.

In summary, this work provides quantitative and qualitative evidence of intracellular MT transfer from MSCs to articular chondrocytes. We demonstrated that inhibition of Cx43 based gap junction signaling prevents MSC-chondrocyte MT transfer. Importantly, we also documented MT transfer to mechanically injured chondrocytes embedded within the native cartilage matrix. This study presents multiple models that may be used to investigate novel biologic approaches to augment MT capacity in chondrocytes and other poorly healing tissues of the skeletal system.

## Materials and Methods

### Primary equine cell co-culture model

Chondrocytes previously harvested from normal femoropatellar joints of young-adult horses (2-5 years) were cultured on T175 plates with Ham’s F12 media (Corning, 1X), with 10% fetal bovine serum (R&D Systems), 1% MEM Amino Acids (gibco, 50X), 2.5% HEPES (Corning, 1M), 1% Penicillin-Streptomycin (Corning, 100X) in 5% CO_2_, 21% O_2_. Bone marrow-derived MSCs were previously harvested from the sternabrae of young-adult equids (2-5 years; n = 3) and cultured on T175 plates with Dulbecco’s Modified Eagle Medium with 1 g/L glucose (Corning, 1X) media with 10% fetal bovine serum (R&D systems), 1% Penicillin-Streptomycin (Corning, 100X), and 0.5% basic fibroblast growth factor (Corning, 1 µg/mL) in 5% CO_2_, 21% O_2_. Passage 2 chondrocytes were plated on a 12-well cell culture plates (250,000 cells per well) for flow cytometry and a 6-well chambered cover glass slide for confocal imaging (60,000 cells per well). At 80% confluence, chondrocytes were stressed by the addition of Rot/A (Sigma Aldrich, 0.5 μM, VWR International, 0.5 μM). After 12 hours of stimulation, chondrocytes were rinsed thrice with 1% PBS, fresh chondrocyte media was replaced, and chondrocytes were stained with 1 μL of cytoplasmic green dye, Calcein AM (Thermofisher, 4 mM). Passage 2 MSCs were stained with MitoTracker Deep Red (Thermofisher, 200 nM), a red dye targeted to intact MT, in a 50 mL conical tube with PBS. The MSCs used in the confocal imaging study were also stained with 5 mg/L Hoechst 33342 (Thermofisher) in the same 50 mL conical tube. The MSCs were incubated in the stain(s) for 30 minutes and rinsed thrice with PBS to prevent residual dyes in the supernatant. The MSCs were then added to wells in a 5:1 chondrocyte:MSC ratio with equal parts chondrocyte and MSC media. The 12-well plates (n = 3) and chambered cover glass slide (n = 1) were incubated for 8 hours. At pre-determined time points (10 minutes, 1, 2, 4, 6, 8 hours), one co-culture well from each of the 12-well plates was lifted with 0.25% Trypsin-EDTA (Corning), fixed with 1% formalin, and stored in PBS at 4°C. At the same time points, 1% formalin was directly added to one well of the 6-well chambered cover glass slide.

### Murine cell co-culture model

Murine experiments were carried out in accordance with animal protocols approved by Institutional Animal Care and Use Committee at Cornell University. UBC mCherry (Jax stock 017614) and PHaM mitoDendra2 (Jax stock 018385) mice were obtained from the Michael Kotlikoff Lab and Serge Libert Labs, respectively, within the East Campus Research Facility at Cornell University. Animals were housed in a Specific Pathogen Free facility. Articular cartilage was harvested from the acetabulofemoral joints of 5-day-old UBC mCherry mice, and chondrocytes were isolated and expanded as previously described (51). Bone marrow-derived MSCs were harvested from 5-week-old PhAM^excised^ mitoDendra2 mice, cultured and expanded as previously described (52) in alpha minimum essential medium (gibco, 1X) with 2.2 g/L sodium bicarbonate, 15% fetal bovine serum, and 1% Penicillin-Streptomycin (Corning, 100X), pH = 7.2. Cell lines were expanded under either hyperoxic (21% O_2_) or physioxic (5% O_2_) conditions from isolation to passage 3, at which cells were used for experiments. Chondrocytes were plated on 12-well cell culture plates (120,000 cells per well), in Dulbecco’s Modified Eagle Medium containing either 1 g/L glucose or 0.45 g/L glucose in 21% O_2_ or 5% O_2_ for 1-3 days. At 75% confluence, chondrocytes were stressed by the addition of either a general inflammatory stimulus, IL-1β (Sino Biological, 10 ng/mL), Rot/A (Sigma Aldrich, 0.5 μM, VWR International, 0.5 μM) or another MT-specific stressor, Oligo (Sigma Aldrich O876, 1 μM) for 12 hours. During stimulation, chondrocyte media did not contain FBS. After 12 hours of stimulation, chondrocytes were rinsed thrice with 1% PBS, fresh chondrocyte media containing FBS was replaced, and passage 3 MSCs were added to each co-culture well in a 10:1 chondrocyte:MSC ratio. After 12 hours of co-culture, cells were lifted with 0.25% Trypsin-EDTA (Corning), fixed in 1% formalin, and then resuspended in PBS with 1% bovine serum albumin.

### Flow cytometry

For both equine and murine cells, flow cytometry was performed using a ThermoFisher Attune NxT analyzer located at the Cornell University Biotechnology Resource Center. Single color controls (non-co-cultured cells) were used to create gates that identified each cell type based on size (using forward and side scatter) and fluorescence. An additional gate was created that identified cells with both green and red fluorescence (red^+^/green^+^ cells), indicating MT transfer. These gates were overlayed onto co-cultured samples in FCS Express (equine) or FlowJo (murine) software.

### Confocal imaging of in vitro co-culture models

Imaging experiments utilized a Leica SP5 confocal microscope located at Cornell University. Formalin-fixed equine cells were imaged on a 6-well chambered cover glass slide (see *Primary equine cell co-culture* model). Live murine cells were imaged longitudinally for up to 9.5 hours after initiation of co-culture in a temperature and humidity-controlled chamber outfitted for the microscope.

### Gap junction inhibition

Murine chondrocytes and MSCs were harvested and expanded as previously described. Chondrocytes were stimulated with Rot/A (Sigma Aldrich, 0.5 μM, VWR International, 0.5 μM) for 12 hours before co-culture. MSCs were treated with either CBX (Sigma Aldrich, 100 μM) or Gap27 (Tocris, 100 μM) for 10 hours before co-culture, and co-culture wells were also treated with each inhibitor for duration of co-culture. Passage 3 MSCs were added to each co-culture well in a 2:1 chondrocyte:MSC ratio. After 24 hours of co-culture, cells were lifted, fixed, and flow cytometry was performed as previously described.

### Chondrocyte gene expression

Chondrocytes were plated on 96 well PCR plates and cultured in physioxia (5% O_2_) or hyperoxia 21% O_2_, in euglycemia (0.45 g/L glucose) or hyperglycemia (1 g/L) and with and without stimulation via Rot/A as above. Briefly, 12-72 hours (at ∼85% confluence) after plating the chondrocytes for experimentation, chondrocytes were either cultured in serum-free media for 12 hours (no stim) or stimulated with Rot/A (0.5 µM/0.5µM) in serum-free media for 12 hours. All wells were aspirated of media, rinsed gently with PBS three times, and then trypsinized to remove cells for RNA isolation. Total RNA was isolated with the RNeasy Mini Kit (Qiagen) according to manufacturer protocol. The High-Capacity cDNA Reverse Transcription Kt (ThermoFisher) was used to generate cDNA according to manufacturer protocol. Sample RNA concentration and quality was determined by Nanodrop. Three reactions (10 uL 2x RT Master mix + 10 uL sample) were performed for each sample. Thermocycler conditions were set at: Step 1-25C, 10 min Step 2-37C, 120 min Step 3-85C, 5 min then Step 4: 4C, until samples were removed. For RT-qPCR, sample cDNA was combined with TaqMan Fast Advanced Master Mix (ThermoFisher) according to manufacturer protocol and added to our custom TaqMan Array Plate. The plates were loaded into the ViiA 7 Real-Time PCR system and a Fast Block 96-well experiment was run according to the ViiA instrument user guide. This experiment was replicated four times (n=4). In the first experiment, technical error in making the hyperglycemic culture medium led to the outliers that were discarded in further analyses.

### Bovine cartilage explant injury and in situ co-culture

Femoral condyle cartilage explants were sterilely harvested from neonatal bovids, obtained post-mortem from a local slaughterhouse, using 6 mm biopsy punches. Explants were cultured in Dulbecco’s Modified Eagle Medium with 0.45 g/L glucose, 2% sodium pyruvate (Corning), 1% L-glutamine (Corning), 1% fetal bovine serum (R&D systems), 2.5% HEPES (Corning, 1M), 1% Penicillin-Streptomycin (Corning, 100X), and 0.5% 1% MEM Amino Acids (gibco, 50X) in 5% CO_2_, 21% O_2_. Explants were subjected to injury using a previously validated spring-loaded rapid-impact model (9). Briefly, the explants were positioned with the articular surface facing up in a PBS filled well and a single, rapid cycle of unconfined axial compression was applied with the impactor. A 10 mm internal spring compression was used, yielding an approximately 17 MPa peak stress and 20 GPa/s peak stress rate as measured by a load cell (50 kHz) at the impactor tip in previous studies. After impacting, explants were immediately returned to media and incubated overnight at 37°C, 5% CO_2_, 21% O_2_ prior to MSC seeding. Prior to seeding, cartilage explants were stained with 100 µM CFDA-SE (Thermofisher) for 30 minutes followed by 30 minutes rinse and incubation in PBS. Passage 5 bovine MSCs previously transduced with mCherry-mito (Vectalys) were seeded onto the articular surface of the assigned explants in bovine MSC media: Dulbecco’s Modified Eagle Medium with 1 g/L glucose (Corning, 1X) media with 10% fetal bovine serum (R&D systems), 1% Penicillin-Streptomycin (Corning, 100X), 0.5% Amphotericin B (Corning, 100X) and 0.5% basic fibroblast growth factor (Corning, 1 µg/mL). First, cartilage media was aspirated from the explant wells and 300,000 MSCs in 15 µl media was pipetted onto the articular surface. For 30 minutes, the 15 µL containing MSCs were incubated on the explants without further addition of cartilage or MSC media to allow maximum seeding. Following this incubation period, cartilage and MSC media was added to the wells in a 1:1 ratio, enough to submerge the explant, and changed every 48 hours for the duration of culturing. Confocal imaging studies were performed 4 days after injury and MSC co-culture. Prior to imaging, bovine explants were bisected into hemicylinders using sterile blades. The rectangular portion of the explants were imaged on a Zeiss LSM880 confocal/multiphoton inverted microscope with 10x and 40x objectives. Images were acquired in two channel sequential scans (green; 488/498-544 and red; 514/563-663 nm excitation/emission, respectively). All parameters were optimized on the first day of imaging, and the same settings were used on subsequent days. Where MSCs were identified, z stacks were obtained (0.75-5 µm spacing in the z plane). Confocal images were captured and imported into Fiji (ImageJ) for further visualization.

### Quantification and Statistical Analysis

Data review and statistical analyses for Fig 1C were performed using GraphPad Prism 9. These data passed the Shapiro-Wilk test for normality (alpha = 0.05). An ordinary one-way ANOVA was performed with post hoc multiple comparisons between group means.

All data review and analyses for Fig 2 (panels C and D) were performed using GraphPad Prism 9 and with consultation from the Cornell Statistical Consulting Unit. Normality was assessed using the Shapiro-Wilk or D’Agostino-Pearson omnibus tests (alpha = 0.05). An ordinary one-way ANOVA was performed with post hoc multiple comparisons between group means.

All data review and analyses for Figure 3 were performed in GraphPad Prism 9. Normality was assessed using the Shapiro-Wilk test (alpha = 0.05). A two-way ANOVA was performed with post hoc multiple comparisons between the control (physiologic conditions) means and the means of hyperoxic and hyperglycemic conditions.

Quantification of gene expression in Figure 4 was done in Excel where delta cycle threshold (ΔCT) was calculated to evaluate relative transcription levels of the sequence of interest compared to the housekeeping gene (GAPDH) and then ΔΔ CT was calculated by subtracting the ΔCT of the gene of interest in experimental conditions from that of control conditions. These data were log transformed in Excel and then further data review and analyses for were performed in Graphpad Prism. Within each gene, equal variance was confirmed by plotting residuals or using the Spearman’s test for heteroscedasticity (P > 0.05). Normality of genes GJA1, TIMP1 and SOD2 was determined using the D’Agostino-Pearson omnibus test and the Shapiro-Wilk test. Then, a two-way ANOVA was performed for each gene.

Differences among group means were considered significant when P < 0.05. Figures were created in GraphPad Prism 9. Sample size (n) represents the number of biological replicates. No statistical methods were used to predetermine sample size.

## Acknowledgments

We thank Lindsay Seewald, Nikita Srivastava, and Rebecca Irwin for their technical support. We acknowledge support by the National Institute of Arthritis and Musculoskeletal and Skin Diseases of the National Institutes of Health (K08AR068470, R03AR075929) and the Harry M. Zweig Fund for Equine Research.

## Data Availability

All study data are included in the article. The gene expression data discussed in this publication have been deposited in NCBI’s Gene Expression Omnibus (53) and will become publicly accessible on September 1, 2022 through GEO Series accession number GSE202788 (https://www.ncbi.nlm.nih.gov/geo/query/acc.cgi?acc=GSE202788).

